# Controllability analysis of macaque structural connectome from an edge centric perspective

**DOI:** 10.1101/2025.03.07.642125

**Authors:** Subham Dey, Eesha Bharti, Zhi-De Deng

## Abstract

In this paper, we investigate the edge controllability properties of the macaque structural connectome, which is reconstructed using optimal tractography parameters. We derive the expression of edge modal controllability and edge average controllability, providing a mathematical framework to analyze their roles from a network systems perspective. Further, we establish the relationship between the two controllability measures, providing insights into their functional implications. We also identify the top edges with the highest average controllability values, which may be critical in facilitating state transitions within the macaque brain network. These findings may have implications for neurostimulation interventions.

## I. INTRODUCTION

White matter tracts facilitate effective information flow between cortical regions throughout the brain [1]. This connectivity of whole brain white matter, often referred to as the structural connectome, supports the brain’s ability to transition between different electrophysiological states, which correspond to different cognitive domains by reflecting neural dynamics associated with processes such as attention, memory, and perception [2]. From the perspectives of control theory [3] and network systems [4], the concept of network controllability can help us to study the dynamical properties of the brain, which are constrained by structural connectome, as well as quantify the energy required to switch between various brain states [5]. Within this framework, two principle measures, average controllability and modal controllability, characterize how easily the brain can move through its energy landscape [6]. Average controllability is related to the energy required for the brain to transition from its current state to a nearby state of less energy. Modal controllability is related to the energy required for the brain to transition from its current state to a state of higher energy. Empirical studies have reported that controllability increases at a rapid rate during the initial infancy [7] to juvenility [8] and well into adulthood [5]. The concepts of network controllability have been exploited to act as a therapeutic planning method for brain stimulation protocols such as electroconvulsive therapy (ECT) [9] and provide a mathematical understanding of neurofeedback and transcranial magnetic stimulation [10]. Moreover, controllability is also known to change significantly in the case of various diverse brain disorders in humans starting from Parkinson’s disease [11], schizophrenia [12], epilepsy [13], and depression [14].

In the human neuroscience literature, the predominant concept of controllability is inspired by the connectivity literature which has a rich history of being a ‘node centric’ phenomenon. In this framework, nodes represent units of information processing, and edges represent entities responsible for transferring information among various network nodes. The enduring popularity of this node centric model stems from its advantages in explaining brain organization, thus strengthening the existing neuroscience literature [15]– [17]. However, the node centric approach has a limitation: it cannot sufficiently explain how systems-level architecture evolves from fluctuations in brain activity, which are inherently time-varying. An alternative approach is an edge-centric model, in which network edges are used to construct the edge connectivity matrix [18]. Nevertheless, most edge connectivity studies focus on data collected while subjects are either in a resting state [19] or engaging in passive viewing tasks, such as movie watching [20]. Contrary to functional connectivity, the literature on edge centric connectivity involving the structural connectome has not been widely explored in the literature [21]. The results related to edge controllability of the structural connectome have also been limited to humans [22]. However, non-human primate models have distinctive similarities to the human brain, cognition, and behavior [23]–[25], therefore it is not unusual for researchers to study the controllability properties of the monkey brain not only from a node centric perspective but also from an edge centric perspective. Non-human primates have a crucial role to play in the field of translational brain stimulation protocols which is supported by research reported in the fields including deep brain stimulation [26], transcranial electric stimulation [27], transcranial magnetic stimulation [28], [29]. As with human research, there are ample opportunities to refine experimental protocols across these domains by integrating network control theory concepts, which motivates the present work.

## II. Mathematical Preliminaries

Control theory dates back to the Industrial Revolution [30]. It concerns the design of control inputs, that is, external perturbations, that guides state transition in a dynamical system, provided that the system is controllable [31] from any initial state to a given desired state. Optimal control theory can be thought of as finding the optimal amount of external perturbations required to make optimal transitions [32]. Network Control Theory (NCT), viewed through the lens of optimal control theory [33], employs mathematical formulations to link the structure and function of the brain [34]. From a node centric perspective, we can model the evolution of brain states over time as a linear time-invariant state-space equation:

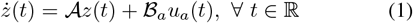

wherein *z* ∈ ℝ^*n*^ represents the brain states, 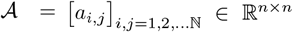 represents the adjacency matrix derived from the structural connectome, ℬ_*a*_ ∈ ℝ^*n×m*^ represents the control input matrix, and *u*_*a*_ ∈ ℝ^*m×*1^ represents the control input to the system. The control input can be considered as external perturbations, such as non-invasive or invasive electromagnetic stimulation to the brain, whereas the control input matrix gives us an idea about how the control input influences the system. The expression given in Eq. (1) can also be written in an equivalent discrete-time setting as

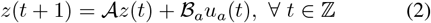

The expression of controllability in the neuroscience literature was initially proposed in the discrete-time domain [5]. The expression of controllability gramian in the discrete-time setting is given by

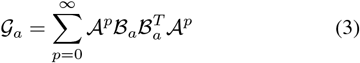

The expression of average controllability is computed as the trace(𝒢_*a*_) in line with previous works reported in the literature as because trace 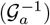 can lead to problems of ill-posed computations [8]. To compute the expression of modal controllability we need the help of eigenvalues of the adjacency matrix 𝒜 [35]. Considering, the matrix 𝒜_*eg*_ = [*r*_*ij*_] which is used to represent the normalized eigenvectors of the adjacency matrix 𝒜. The term *r*_*ij*_ reflects the controllability of the mode *λ*_*j*_(𝒜) provided the control node is the node i. Thus, with the help of Popov Belevitch Hautus (PBH) test [36] we can express the scaled measure of controllability of all modes 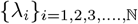 from the control node *i* as

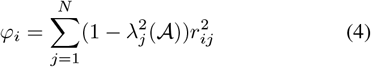

Now, we are in a situation to have an expression of the adjacency matrix of the network system from an edge centric perspective but to do so we need the expression of incidence matrix which is given by [37]

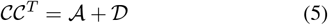

where 𝒟 = diag(*d*_*ii*_)_*i*=1,2,…,N_ and *d*_*ii*_ = Σ_*j*_ (*a*_*ij*_). The expression of the edge adjacency matrix is given by

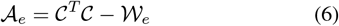

where 𝒲_*e*_ represents a matrix that is diagonal, with each of its elements representing the weight of each edge. A pictorial representation of the concept related to edge line graph and its original graph is given in Fig. 1 for better visual representation, where the number of nodes is 6. In the line graph representation, each edge of the original graph becomes a node in the line graph.

**Fig. 1.**
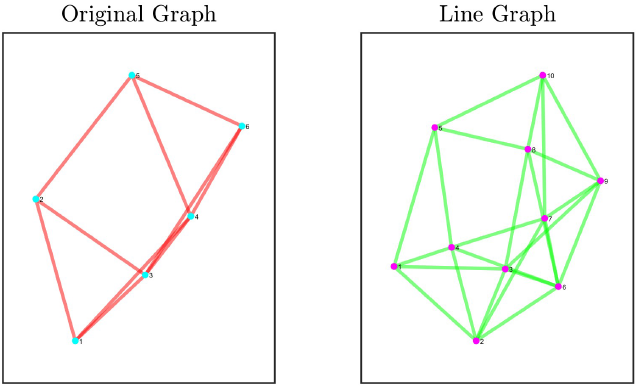
Original graph and its corresponding edge graph wherein the nodes are numerically numbered for the original graph whereas the edges are numerically numbered in case of the line graph.

Similar to the node centric average controllability, the expression of edge centric average controllability can be obtained with the help of a gramian matrix. If the control matrix to the edge centric line graph is given by ℬ_*e*_, then the corresponding gramian matrix is given by

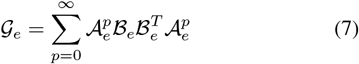

The expression of edge average controllability is given by trace (𝒢_*e*_). Similarly, the expression of edge modal controllability is given by

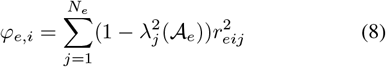

The term *N*_*e*_ is used to represent the dimension of the edge adjacency matrix 𝒜_*e*_. The term *r*_*eij*_ represents the normalized eigenvectors of the edge adjacency matrix. A pictorial representation of the energy landscape which can be used to explain the concepts of edge modal controllability and edge average controllability is given in Fig. 2.

**Fig. 2.**
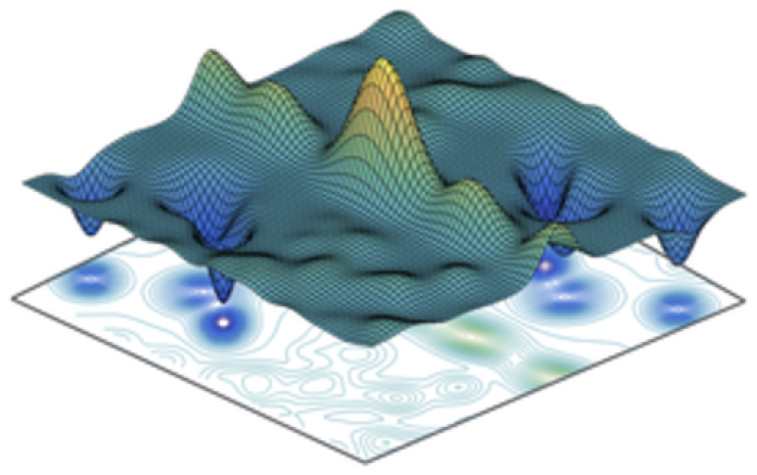
A pictorial representation of the energy landscape plot wherein the peaks represent the higher value of energy whereas the troughs represent the lower value of energy. An edge is said to have a high value of edge average controllability if stimulating that edge leads to a transition from global minima of the energy landscape to a nearby local minima whereas an edge is said to have a high value of edge modal controllability if stimulating that edge leads to a transition from the global minima to global maxima of the energy landscape.

## III. Dataset

Macaque structural connectome can be constructed using various methods. One method relies on neuroanatomical data obtained from tract-tracing databases, such as the CoCo-Mac database [38], [39]. Another popular tracer dataset is provided by Markov et al. [40], which offer a weighted description of the macaque connectome. Yet, one significant limitation of the method proposed in [40] is that it only has 29 of the 91 regions of the macaque single hemisphere represented in it. An alternative approach to constructing the connectome is through the use of the tractography method on diffusion weighted imaging dataset similar to what we see in the human counterparts [41]. The probabilistic tractography method plays a satisfactory role in finding fiber tract capacities and tract length in macaques [42]–[44]. However, a limitation of probabilistic tractography is that it is susceptible to false positive connections, which can significantly impact the reconstructed network topology [45]–[48]. The most comprehensive reconstructions emerge by combining both tracer data and tractography methods [49].

Previous research [50]–[52] demonstrates that tractography parameters critically affect its accuracy. Leveraging on the insights obtained from the previous studies, the tractography algorithm used in this paper is first optimized to replicate the best weighted partial cortex tracer-based connectome as reported in [40] and then extending it to obtain full cortex connectome weights. Also, the tract lengths can be obtained from the tractography algorithm, which for tracer datasets can only be calculated in terms of geodesic or Euclidean distances of regions of interest (ROIs). The tract lengths are important if one wants to perform whole brain simulations.

In our work, we used two different datasets available in the literature. First, the tracer derived connectivity matrix as described in [53] with Kötter and Wanke parcellation [54] was derived from the CoCoMac dataset [38], [39] with 82 ROIs. The second dataset is reported in [40]. The average connectome matrix contains contributions from a total of nine subjects (eight *Macaca mulatta* and one *Macaca fascicularis*) [42]. Experimental protocols involving image acquisition and preprocessing are mentioned in the previous papers [44], [49], [55]. The atlas used for connectome construction is the F99 macaque atlas [56]. FSL software was used to perform tractography [57]. Various curvature thresholds with steps of 0.2 between 0.2 and 0.8 as well as the effects of distance correction were taken into consideration simultaneously to find out the optimal parameter values which gives rise to the most correct connectome with the 29 ROI dataset as the reference. Once the optimal set of parameters were obtained, it was used to perform tractography on the Regional Map (RM) parcellation using the F99 atlas. Final undirected and symmetric connectome was a result of average of tractography outputs for each of the subjects. We use this averaged connectome for proving our theoretical concept of edge controllability. However, similar results can be trivially obtained for each subject individually.

## IV. Results

The node adjacency matrix of the macaque structural connectome is a square matrix of dimension 82. For an undirected graph, the edge adjacency matrix has a dimension of 3321 (calculated as 0.5 × *n* × (*n* − 1), where *n* is the number of nodes). Past research has shown that node modal and node average controllability have an inverse relationship: when modal controllability is high, average controllability is low, and vice versa. This follows directly from the energy landscape framework, which forms the backbone of the controllability analysis. Extending this idea, we observe in Fig. 3, a high anti-correlation measured using the Pearson correlation coefficient measure (*r* = − 0.9967, *p <* 10^*−*5^) between edge modal controllability and edge average controllability. Fig. 3 also suggests that the distribution of average controllability values is skewed, with relatively few edges exhibiting high average controllability and the majority falling at lower values. Based on RM parcellation some of the top edges of high values of average controllability are given in Table I. The numerical values of the controllability metrics are of not much importance because controllability is a relative measure and does not play a role in calculating control input, which results in optimal interventions.

**Fig. 3.**
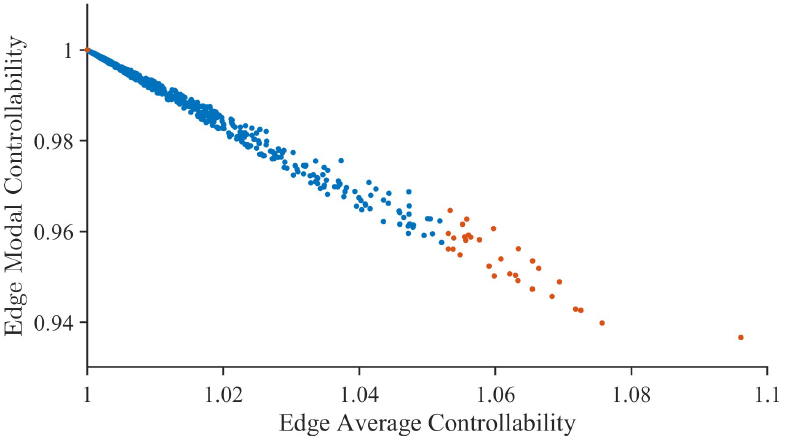
Relationship between edge average controllability and edge modal controllability for the macaque structural connectome, with the top 30 and bottom 30 values of edge average controllability highlighted in red.

**TABLE I.** Edges having highest average controllability values.

| Edge | Edge average controllability value |
| --- | --- |
| Left-right anterior cingulate ctx | 1.0961 |
| Left gustatory ctx-left ventrolateral premotor ctx | 1.0757 |
| Right medial parietal ctx-right intraparietal ctx | 1.0726 |
| Right posterior cingulate ctx-right medial parietal ctx | 1.0718 |
| Left ventrolateral prefrontal ctx-left centrolateral prefrontal ctx | 1.0694 |
| Left superior temporal ctx-left primary auditory ctx | 1.0683 |
| Right anterior cingulate ctx-right medial premotor ctx | 1.0664 |
| Left anterior cingulate ctx-left medial premotor ctx | 1.0655 |
| Left posterior cingulate ctx-left medial parietal ctx | 1.0654 |
| Left ventral temporal ctx-left centrolateral prefrontal ctx | 1.0634 |

ctx: cortex

## V. Discussions

In this work, we examined the controllability properties of the macaque structural connectome from edge centric perspectives. Our focus on the edge adjacency matrix and associated controllability metrics, rather than confining the analysis to nodes, was motivated by growing evidence that inter regional connections (i.e., white matter tracts) offer unique insights into how brain dynamics can be influenced or controlled [58]. The results confirm and extend previous findings in human datasets [22] by demonstrating a marked inverse relationship between edge average controllability and edge modal controllability (Fig. 3). This reflects the idea that network features are well-suited for low-energy transitions in brain states (average controllability) tend not to coincide with those enabling high energy transitions (modal controllability).

A second key finding lies in the distribution of edge controllability values. Our data reveal a skewed distribution for edge average controllability, with fewer edges having high values and the majority having low values. This skewness has significant implications for both theoretical modeling and experimental design. Edges with exceptionally high average controllability (Table I) may represent prime targets for interventions aimed at smoothly guiding the brain toward functionally relevant state. Whether these states pertain to improved mood in depression, modulation of seizure thresholds in epilepsy, or enhanced cognitive performance, the targeted edges could be sites of stimulation for invasive or noninvasive modalities. In contrast, edges with higher modal controllability might be better suited for abrupt state changes potentially useful in interrupting pathological network dynamics.

These observations highlight the importance of considering white matter tracts as key agents of causal control, echoing recent developments seen in human neuromodulation research. Both invasive and noninvasive brain stimulation approaches are shifting from node focused interventions to those that target inter regional connections. In neurofeedback, for example, researchers increasingly analyze edge-level connectivity in human subjects [59]. Similarly, for deep brain stimulation (DBS), there has been a paradigm shift [58] toward targeting white matter tracts rather than strictly localized anatomical regions, as illustrated by better treatment responses in patients with treatment-resistant depression when targeting the convergence of forceps minor, cingulum bundle, and subcortical-junction white matter pathways [60]. A parallel shift can be seen in transcranial magnetic stimulation (TMS). Traditionally aimed at the left dorsolateral prefrontal cortex (DLPFC), TMS has proven effective in treating depression, yet individual patient outcomes vary greatly. Recent work has shown that connectivity-based targeting particularly the functional connectivity between the DLPFC and remote regions such as the subgenual cingulate is key to optimizing TMS efficacy [61]. Collectively, these efforts underscore that the future of brain stimulation may hinge on connection level approaches, aligning with our findings that edges serve as powerful agents in network level control.

Historically, nonhuman primates, particularly in the context of Parkinson’s disease and DBS have proven indispensable for refining stimulation protocols [62]. Our work adds a layer of precision to brain stimulation experiments by stimulating edges with high values of average controllability [62]. Similarly, our research can help design optimal experimental paradigms for transcranial direct current stimulation [63], transcranial electric stimulation [27], and transcranial magnetic stimulation [28], [29] for simian species. Edge controllability analysis has the added advantage of more information compared to their node controllability counterparts [9].

Several limitations warrant further attention. First, although we derived our conclusions from carefully curated tractography protocols and tracer-based connectivity, the methods for constructing edge level network representations vary across datasets and can introduce bias. Second, our averaged connectome provides characterization of group features, but real world applications particularly therapeutic interventions may require individualized mapping to accommodate the unique structural features of each subject. Finally, while we emphasize controllability in a linear approximation, the true dynamics of neural systems may deviate from linearity during certain tasks or pathological states [10]. Future investigations that adapt edge controllability concepts to nonlinear or time-varying frameworks could yield deeper insights.

## VI. Conclusions

In summary, our work explores the application of edge centric controllability method to macaque structural connectome, which can be used to improve the experimental efficacy of a wide range of brain stimulation paradigms in simian species and therefore play a pivotal role in the field of translational brain research.

## Acknowledgment

The authors are supported by the National Institute of Mental Health (NIMH) Intramural Research Program (ZI-AMH002955). This work utilized the computational resources of the NIH HPC Biowulf cluster (http://hpc.nih.gov).

## Notes

### Competing Interest Statement

The authors have declared no competing interest.

### Summary of Updates

I have modified some text and corrected some typos in the manuscript.

